# Viscosity Prediction of High-Concentration Antibody Solutions with Atomistic Simulations

**DOI:** 10.1101/2023.08.28.555069

**Authors:** Tobias M. Prass, Patrick Garidel, Michaela Blech, Lars V. Schäfer

## Abstract

The computational prediction of the viscosity of dense protein solutions is highly desirable, for example in the early development phase of high-concentration biopharmaceutical formulations where the material needed for experimental determination is typically limited. Here, we use large-scale atomistic molecular dynamics (MD) simulations with explicit solvent to *de novo* predict the dynamic viscosities of solutions of a monoclonal IgG1 antibody (mAb) from the pressure fluctuations using a Green-Kubo approach. The viscosities at simulated mAb concentrations of 200 mg/ml and 250 mg/ml are compared to the experimental values, which we measured with rotational rheometry. The computational viscosity of 24 mPa s at a mAb concentration of 250 mg/ml matches the experimental value of 23 mPa s obtained at a concentration of 213 mg/ml, indicating slightly different effective concentrations (or activities) in the MD simulations and in the experiments. This difference is assigned to a slight underestimation of the effective mAb-mAb interactions in the simulations, leading to a too loose dynamic mAb network that governs the viscosity. Taken together, the present study demonstrates the feasibility of all-atom MD simulations for predicting the properties of dense mAb solutions and provides detailed microscopic insights into the underlying molecular interactions. At the same time, it also shows that there is room for further improvements and highlights challenges, such as the massive sampling required for computing collective properties of dense biomolecular solutions in the high-viscosity regime with reasonable statistical precision.

**TOC Graphic:** 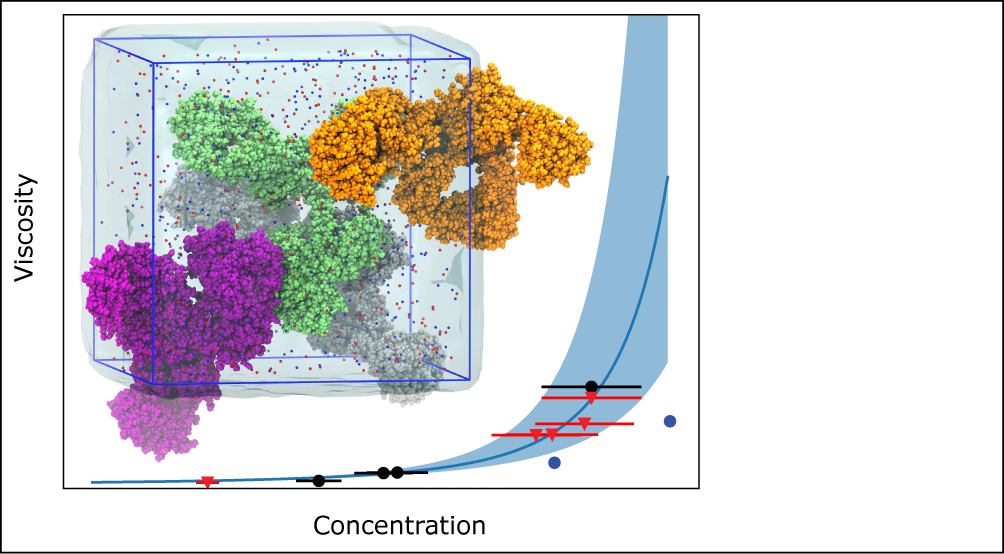

## Introduction

Highly concentrated solutions of biomolecules play an important role in both biology and pharmaceutical applications. Up to thirty percent of the interior volume of biological cells is filled with macromolecules, and such densely packed, “crowded” environment can have an influence on biomolecular structure and thermodynamic stability, as well as on the kinetics of intracellular processes.^1–5^ Likewise, for biopharmaceutical applications, reaching high protein concentrations as required for subcutaneous drug dosages, the crowded solution faces different challenges such as colloidal protein stability, the formation of protein particles, phase separation, and increased viscosity.^6^ Such formulation aspects are particularly challenging for large and conformationally flexible molecules, such as monoclonal antibodies (mAbs). MAbs are one of the most important classes of biopharmaceuticals, with mAb-based immunotherapies having found widespread applications ranging from the treatment of cancer^7^ to COVID-19.^8^ Subcutaneous injection is a preferred route of administration. However, the high dosages required (ca. 1 mg of mAb per kg of body weight) and the small volumes that can be injected under the skin or into the muscle (less than 2 ml) imply high-concentration formulations, typically higher than 100 mg/ml.^9^ The injection needles used for subcutaneous application are usually thin for patient convenience (injection needles with 27 G to 29 G for subcutaneous injection and even thinner for intravitreal injection with 30 G to 32 G), and a high viscosity of the formulation is especially problematic as it is connected to high injection forces and tissue back pressure. Furthermore, high viscosity has been linked to the irreversible formation of protein particles, which reduces the efficacy of the drug and can cause unwanted immunogenicity.^10^

At the molecular level, macroscopic collective properties of high-concentration protein solutions are governed by protein-protein interactions, which lead to the transient formation of dynamic, nonspecific clusters.^11–13^ In principle, molecular dynamics (MD) simulations are a powerful tool to predict the properties of concentrated protein solutions because solely basic physical laws and established physicochemical principles are used. Molecular simu-lations have been used to investigate the structure and dynamics of the crowded cellular cytoplasm, both using molecular-level Brownian dynamics (BD) simulations with implicit solvation models,^14,15^ coarse-grained (CG) models,^16^ and also at the fully atomistic level,^17,18^ see Ostrowska et al. ^19^ for a recent review. All-atom and coarse-grained MD simulations have also been used to characterize biomolecular condensates, as for example formed by liquidliquid phase separation.^20–23^ Simulations of dense antibody solutions are typically performed with colloidal hard-sphere models^24–26^ or with super-CG models in which each individual domain of the mAb is represented by a single CG bead, resulting in the representation of the entire mAb by only a dozen CG beads.^27–30^ Such models are computationally very cheap and thus allow one to simulate large systems with a large copy number of mAbs in the simulation box, and to screen a range of different conditions in the simulations such as mAb concentration, ionic strength of the solution, etc. However, the approximations inherent to the CG models can limit their accuracy and typically require extensive parametrization and calibration against experiments and/or higher-level computations, thus hampering the predictive capacity of such CG models. To our knowledge, no *de novo* predictions of dynamic viscosities of concentrated mAb solutions have been reported on the basis of such CG simulations.

The desired accuracy can be achieved with all-atom MD simulations with explicit solvent, which automatically also take excluded volume effects, (solvent-mediated) short- and long-range interactions, and hydrodynamic interactions into account. All-atom MD should therefore be truly predictive, in the sense that no experimentally-derived adjustable parameters are needed. However, due to the huge computational costs associated with fully atomistic simulations of such large and flexible molecules, all-atom MD studies of complete mAbs were so far limited to single molecules^31–35^ or dimers.^36^

Concerning the viscosity of such concentrated biomolecular solutions, several MD-based methods have been developed for determining the shear (dynamic) viscosity of a liquid, both with equilibrium as well as nonequilibrium simulations.^37^ One appealing approach is the Green-Kubo (GK) integral, which relates the viscosity *η* to an integral over the pressure tensor time autocorrelation function (ACF),^38^

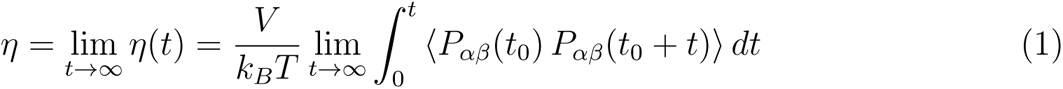

where *V* is the volume of the simulation box, *k_B_* is Boltzmann’s constant, and *T* is the temperature. The angular brackets ⟨*…*⟩ indicate thermal averaging. *P_αβ_* are the elements of the pressure tensor, whose ACFs are calculated by averaging over all time origins *t*_0_. The pressure tensor can be readily obtained from an equilibrium MD simulation. In principle, equation 1 enables a direct evaluation of the viscosity from the asymptotic limit of the GK integral, without the need to invoke a model such as Stokes-Einstein. However, the pressure fluctuations in a microscopic (nm-sized) system are very large, rendering the accurate estimation of *η* challenging due to slow convergence of the GK integral at long correlation times. The accumulation of noise in the tail of the ACF, which slowly decays to zero, can lead to a large degree of variability in the running integral.^37^ Formally, equation 1 needs to be integrated to infinity. In practice, the integral is carried out until *η*(*t*) plateaus at a stable value, which can be difficult to unambiguously identify due to the fluctuations (see below).

Recently, von Bülow et al. conducted large-scale all-atom MD simulations in explicit solvent to characterize the diffusional dynamics and viscosity of dense protein solutions.^39^ The highest protein concentrations studied were 200 mg/ml, with simulation system sizes ranging up to 3.6 million atoms. This impressive work demonstrated the strong slowdown of translational and rotational protein diffusion with increasing concentration, which can be quantitatively explained with a dynamic cluster model.^39^ The study provides deep insights into the molecular interactions and dynamics that are at play in dense protein solutions. However, due to the huge computational costs, the study of von Bülow et al. focused on small and rigid model proteins, such as ubiquitin (M*_W_* ca. 8.5 kDa), hen egg white lysozyme (ca. 14.3 kDa), and villin headpiece (ca. 7.3 kDa), and it was limited to the low-viscosity regime (*η <* 3 mPa·s).^39^

Here, we explore the application of large-scale atomistic MD simulations with explicit solvent for highly concentrated and viscous solutions of large and conformationally flexible proteins, such as mAbs. As a real-life application, we selected the humanized monoclonal IgG1 antibody of Padlan^40^ (M*_W_* ca. 150 kDa) as a model system (Figure 1A). All-atom MD simulations of four complete antibody molecules in the simulation box (Figure 1B) were conducted at mAb concentrations of 200 mg/ml and 250 mg/ml; each of the two systems was simulated for a total time of 7.8 *µ*s. The simulated mAb concentrations are similar to the concentrations of 199 and 213 mg/ml at which we experimentally determined the dynamic viscosity of the mAb using rotational rheometry. The shear viscosity of the mAb solutions was computed from the pressure fluctuations during equilibrium MD simulation via the Green-Kubo integral (equation 1). The computational results are compared to and critically discussed in light of the experimental viscosity.

**Figure 1:**
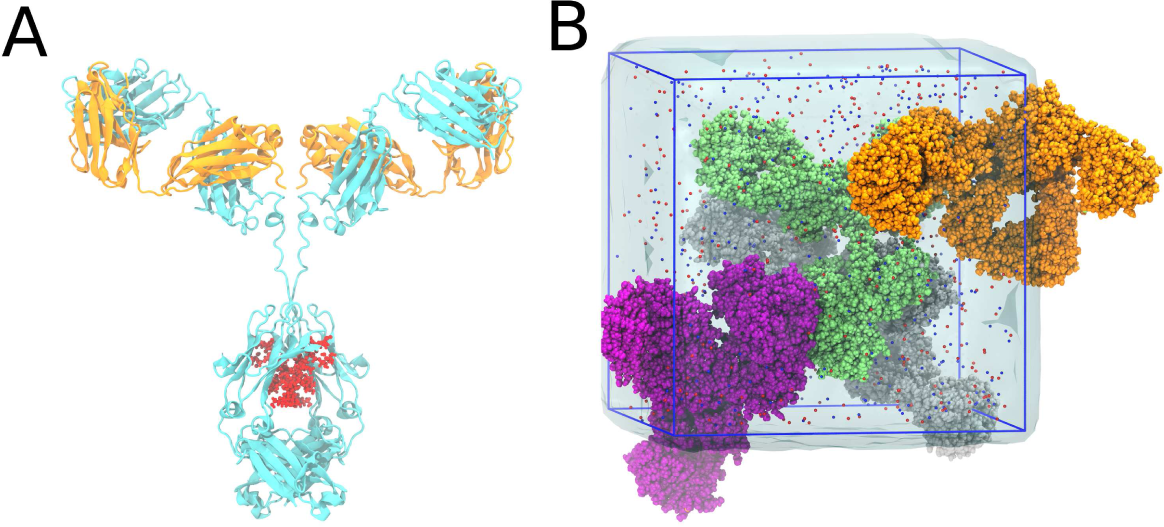
A) The Padlan IgG1 mAb is shown in cartoon representation, with light and heavy chains colored in orange and cyan, respectively. The glycan in the Fc domain is shown as red spheres. B) MD simulation system comprised of four mAbs solvated by water and 150 mM NaCl.

The rest of the manuscript is organized as follows. In the Methods section, the details of the MD simulations and their analysis in terms of the time decomposition approach suggested by Zhang et al. ^41^ are described, which aims at a reliable estimation of the viscosity from the long-time limit of the GK integrals. Furthermore, the rheometry experiments are explained. In the Results section, we demonstrate the application of the approach for the present case. The viscosities predicted from the computations are analyzed with a particular focus on the statistical errors, and are compared to the experimentally determined viscosity. In the Discussion section, we consider possible issues on the computational side, especially those related to the force field and to the limited time scale of the MD simulations and the slow convergence of the GK integrals. In addition, also possible limitations of the rheometry experiments are critically discussed, which render the comparison of simulated and measured viscosities challenging, at least at a quantitative level. The final Conclusions section summarizes the main points of the work and briefly outlines the possible relevance of the present findings for other, related applications, such as molecular simulations of biomolecular condensates and of the crowded cytoplasm of biological cells.

## Methods

### Computational Methods

#### Simulation Setup

All simulations were carried out with the GROMACS code (version 2020.5).^42^ The atomic coordinates of the Padlan mAb were taken from http://www.umass.edu/microbio/rasmol/ padlan.htm. The AMBER a99SB-disp^43^ force field was used for the protein and the water, and the glycan moieties were described with the GLYCAM06j force field.^44^ To set up the simulation systems, four mAb molecules were placed randomly inside a cubic simulation box with a volume corresponding to the desired target concentration (edge lengths of ca. 17.0 and 15.8 nm for *ρ* = 200 and 250 mg/ml, respectively). The random initial placement of the individual mAbs was accepted or rejected based on atomic clashes, and repeated until all mAb molecules were accommodated in the box. The titratable amino acid side chains were protonated according to their standard charge states at pH 7. The boxes were filled with water, and the total charge of the simulation box was neutralized by adding sodium and chloride ions at a concentration of 150 mM, yielding final system sizes of ca. 633.000 and 509.000 atoms for 200 and 250 mg/ml, respectively. This random placement procedure was repeated three times to generate an independent starting configuration for each of the three repeats. The large sizes of the simulation systems and the very long sampling times needed for the viscosity calculations render the study of finite box-size effects prohibitively expensive, as already simulation boxes with 8 mAbs will comprise of more than 1 million atoms.

The applied protocol to set up the simulations is expected to be advantageous over using the same starting coordinates and merely generating different random atomic velocities to initialize the MD simulations, because different mutual orientations of the mAbs in the box are constructed. The rearrangement of the mAbs is very slow and cannot be expected to be fully sampled during the MD simulations.

#### MD Simulations

The simulations were carried out in cubic simulation boxes, with periodic boundary conditions. Prior to the production simulations, the systems were energy minimized (2000 steps of steepest descent) and equilibrated for 2 ns at 298 K and 1 bar with harmonic position restraints on all mAb heavy atoms (force constants of 1000 kJ mol*^−^*^1^ nm*^−^*^2^). Virtual site hydrogens were used for the protein together with hydrogen mass repartitioning (HMR) between the H-atoms and heavy atoms of the glycan, allowing to integrate the equations of motion with 4 fs time steps. Bond lengths were constrained using the LINCS algorithm.^45^ The number of matrices in the expansion for matrix inversion was set to 6 (“LINCS-order” parameter in GROMACS) to more accurately describe the contributions of the constraint forces to the virial and thus to the pressure. SETTLE^46^ was used to constrain all internal degrees of freedom of the water molecules. Constant temperature (298 K) and pressure (1 bar) during the simulations were ensured via the Bussi velocity rescaling thermostat^47^ and the Parrinello-Rahman barostat,^48^ respectively, with coupling time constants of *τ_T_* = 0.5 ps and *τ_p_* = 10 ps (a Berendsen barostat was used during the first 300 ps of the 2 ns position-restrained equilibration). A buffered Verlet neighbor list^49^ was used for the pairwise non-bonded interactions. Lennard-Jones 6,12 interactions were smoothly shifted to zero at 1.0 nm distance cutoff. An analytical correction was added to the energy and pressure to correct for this truncation of the Lennard-Jones interactions. The 1.0 nm distance was also used to switch between the short-range and the long-range Coulomb interactions, which were treated with the smooth particle-mesh Ewald (PME) method^50,51^ with a 0.12 nm grid spacing and cubic B-spline interpolation.

For each of the two mAb concentrations investigated (200 and 250 mg/ml), three production simulations of length 500 ns were carried out. From these three repeats, the first 100 ns were considered to be equilibration and discarded. From the last 400 ns, 21 snapshots were extracted (evenly spaced in time, that is, after 100, 120*, …,* 500 ns) and used as starting configurations for 100 ns simulations at constant volume (in the canonical ensemble). During these NVT simulations, the pressure tensor was stored to disk every 4 fs to capture the fast fluctuations (fast oscillations in the pressure ACF).

### Viscosity Computation

The nondiagonal elements of the pressure tensor, *P_xy_*, *P_yz_*, and *P_xz_*, were recorded. In addition, also the following combinations of diagonal elements were recorded to improve statistics:^52,53^ (*P_xx_* − *P_yy_*)*/*2, (*P_xx_* − *P_zz_*)*/*2, and (*P_yy_*− *P_zz_*)*/*2. All pressure-pressure ACFs were computed up to a correlation time of 20 ns according to equation 1 and averaged, after already averaging over the corresponding pairs *P_αβ_, P_βα_* (for example *P_xy_* and *P_yx_*). The ACFs from all 21 NVT runs that were spawned from each of the three 500 ns repeat simulations were averaged. The separate analysis of the three repeats was used to provide a conservative estimate of the statistical uncertainties. Finally, the data from all 63 NVT runs were averaged.

To compute the viscosities from the long-time limits of the GK integrals, the time de-composition method suggested by Zhang et al. ^41^ was employed. The procedure is as follows. For each of the *N* = 63 individual MD trajectories generated for each of the two mAb concentrations, *η*(*t*) was calculated based on the GK relation (equation 1). These individual GK running integrals were averaged over the *N* trajectories to obtain ⟨*η*(*t*)⟩ and the corresponding standard deviation

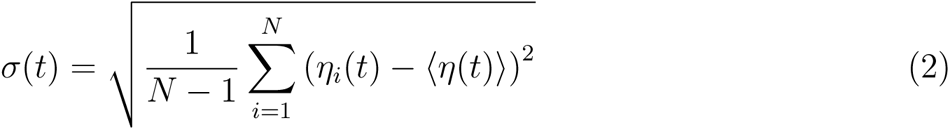

which was fitted with the power law

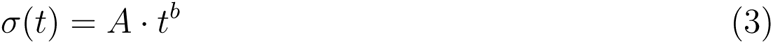

with the parameters *A, b >* 0. To eliminate the noise in the GK integrals, ⟨*η*(*t*)⟩ was fitted by the triexponential function

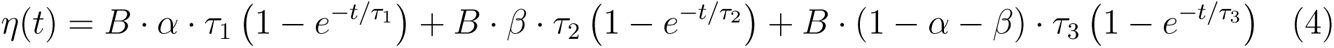

with fitting parameters *B >* 0, *α, β <* 1, and *τ*_1_,_2_ *>* 0. In this fit, the data were weighted with *t^−b^*, with the exponent *b* determined from the power law fit (equation 3). All data are used for the fit, but the statistically accurate data at short *t* have larger weights in the fit than the more noisy data at longer times (according to *σ*(*t*)), thereby minimizing the uncertainty of the estimated viscosity. In this way, the nonuniform weighting implements a tradeoff between statistical errors on the one hand (which favor short *t*) and systematic errors on the other hand (which impose long *t*, equation 1). By construction, the analytic function asymptotically converges, and the final estimates of the viscosity values are taken as the long-time limit of equation 4.

Equation 4 results from integrating an ACF that is described by a triexponential decay. However, we consider the choice of a triexponential function to be purely empirical (in fact, the asymptotic decay of the pressure ACF is rather *t^−^*^3/2^),^37^ and although the time constants *τ* might be associated with different dynamic processes in the solution, we do not assign a physical interpretation to the parameters. In their studies of more simple liquids, Hess ^37^ and Rey-Castro and Vega ^52^ used biexponential functions. For the present case, we found that biexponential functions cannot capture the slow rise of *η*(*t*) and thus do not describe the actual data as well as the triexponentials, see Figure S1 in Supporting Information. Such long transients in the running integrals of *η*(*t*) are absent for the more simple liquids investigated by Hess ^37^ and Rey-Castro and Vega.^52^

#### Bootstrapping analysis

A bootstrapping approach was followed to further investigate the statistical convergence of the computed viscosity values. From the full set of 63 individual viscosity curves (GK running integrals), a subset of curves was repeatedly drawn randomly (without replacement, that is, every trajectory was only drawn once for each separate draw). Subsets of increasing size ranging from 10 to 60 trajectories were drawn 100 times, averaged, and fitted with the triexponential function to evaluate the viscosity in the long-time limit (equation 4). The average and the standard deviation of the viscosities obtained this way were determined.

### Experimental Methods

#### Monoclonal Antibody Formulation

The Padlan monoclonal antibody investigated in this study was produced by mammalian cell culture technology using Chinese hamster ovary (CHO) cells.^54,55^ The mAb was purified using affinity and ion-exchange chromatography according to Jacobi et al. ^55^ A complete buffer exchange was achieved by extensive tangential flow filtration, according to Bahrenburg et al.,^56^ followed by dialysis carried out in a Slider-A-Lyzer dialysis cassette (Thermo Scientific, Rockford, USA) against 10 mM histidine, 150 mM sodium chloride, pH 7 (formulation buffer). The solution was concentrated to 200 mg/ml by Amicon Ultra-15 Centrifugal Filter Units with a molecular weight cutoff of 30 kDa at 25 °C and 5′000 g. An aliquot was filtrated through sterile Sterivex-GV Filter units (Millipore Corporation, Billerica, USA) whereas the other remained unfiltered. Lower concentrated mAb solutions were produced by diluting the 200 mg/ml mAb solution with the corresponding formulation buffer. Sample quality was ensured by quantification of high molecular weight (HMW) and low molecular weight (LMW) species using analytical size exclusion chromatography (SEC).

#### Dynamic Viscosity

Dynamic viscosity was measured using a HAAKE Mars III rotational rheometer (Thermo Scientific) equipped with a 35 mm titanium cone (cone angle 1*^◦^*). A volume of 200 µl of mAb sample was pipetted on a static surface and equilibrated for 5 minutes. The generated shear forces were analyzed via torque measurements. The relation between dynamic viscosity (η) torque (*M*), angular velocity (ω), cone angle (α), and cone radius (*R*) is

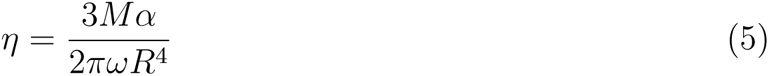

according to Hartl et al. ^57^ and Mezger.^58^ The viscosity was measured in controlled shear rate (CSR) mode with a rotation ramp of *τ* = 100−1000 s*^−^*^1^ in 10 logarithmic steps. The dynamic viscosity was measured at a constant temperature of 297 K and a continuous shear rate of *τ* = 1000 s*^−^*^1^ for 100 s, with measurements averaged over 1 s intervals. Afterwards, the shear stress was recorded on a temperature ramp from 250 K to 288 K in single measurements at a shear rate of 1000 s*^−^*^1^. The data were analyzed using the Haake RheoWin Data Manager software (Thermo Fisher Scientific). The viscosity data at 297 K was used to fit the Ross-Minton equation^59^

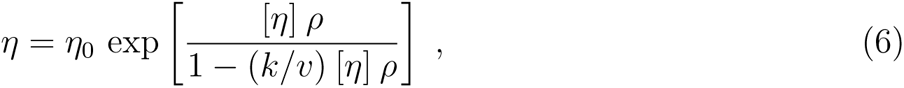

where *η*_0_ is the viscosity of the buffer solution, [*η*] is the intrinsic viscosity of the solute, *c* is the mass concentration, *k* is a crowding factor and *v* is a parameter that corresponds to the shape of the solute molecule (*v* = 2.5 for a perfect sphere, *v >* 2.5 for non-spherical particles).

## Results

The Green-Kubo formula (equation 1) involves the integration over the pressure-pressure autocorrelation functions (ACFs), which are plotted in Figure 2 for the two mAb concentrations investigated. The features of the ACF at short correlation times (Figure 2, inset) differ in a distinct manner from those of bulk a99SB-disp water, see Figure S2 in Supporting Information for a comparison. The ACFs have a long tail, which slowly decays to zero (note the logarithmic scale in the plot). This decay is somewhat slower at 250 mg/ml mAb concentration than at 200 mg/ml, in line with the expectation that the dynamics are sluggish at higher protein concentration.

**Figure 2:**
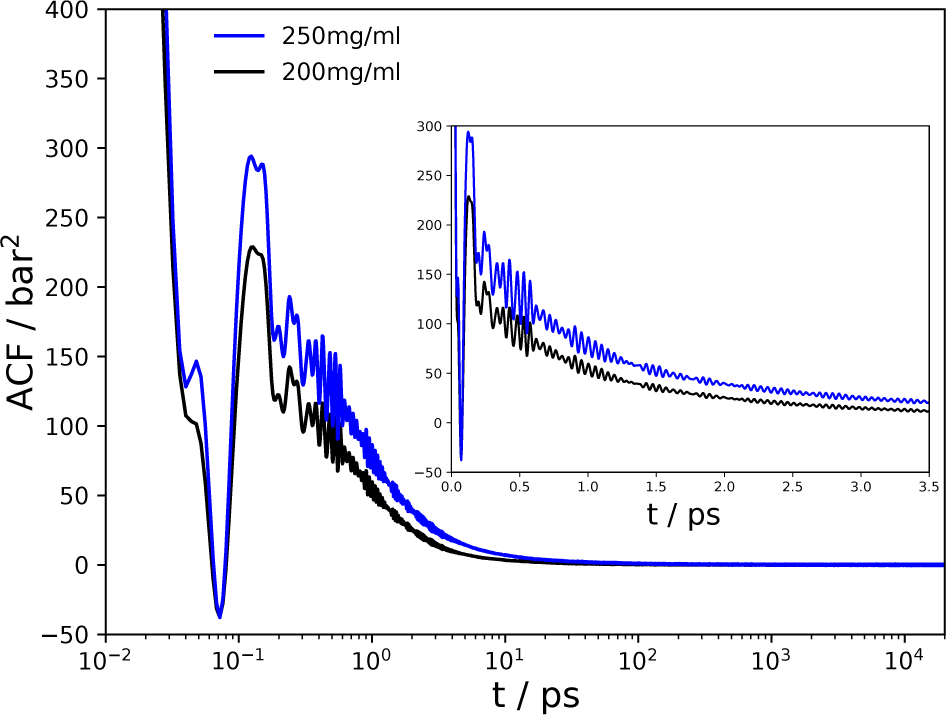
Pressure ACFs for the mAb solutions at 200 mg/ml (black) and 250 mg/ml (blue) concentrations. The ACFs shown were averaged over all nondiagonal elements of the pressure tensor as well as over the contributing combinations of diagonal elements (see Methods), and also averaged over all 63 individual trajectories. The inset shows the short-time behavior of the ACFs (linear time axis).

To determine the shear viscosities from the long-time limits of the GK integrals (equation 1), the method suggested by Zhang et al. ^41^ was used, which minimizes uncertainties due to the increasing long-time noise of the ACFs and the resulting fluctuations of the GK integrals (see Methods). Figures 3A and B show the GK integrals from the 63 individual MD trajectories for the mAb concentrations of 200 and 250 mg/ml, respectively (thin grey lines). The average curves are shown as thick lines, together with the triexponential fits (equation 4). The substantial noise in the individual curves is very much reduced in the averages (Figure 3C), and the fits yield robust viscosity estimates in the long-time limit. The viscosities predicted by the MD simulations are 9.2 ± 5.8 mPa·s and 24.1 ± 7.8 mPa·s for the mAb solutions at concentrations of 200 mg/ml and 250 mg/ml, respectively (Table 1).

**Figure 3:**
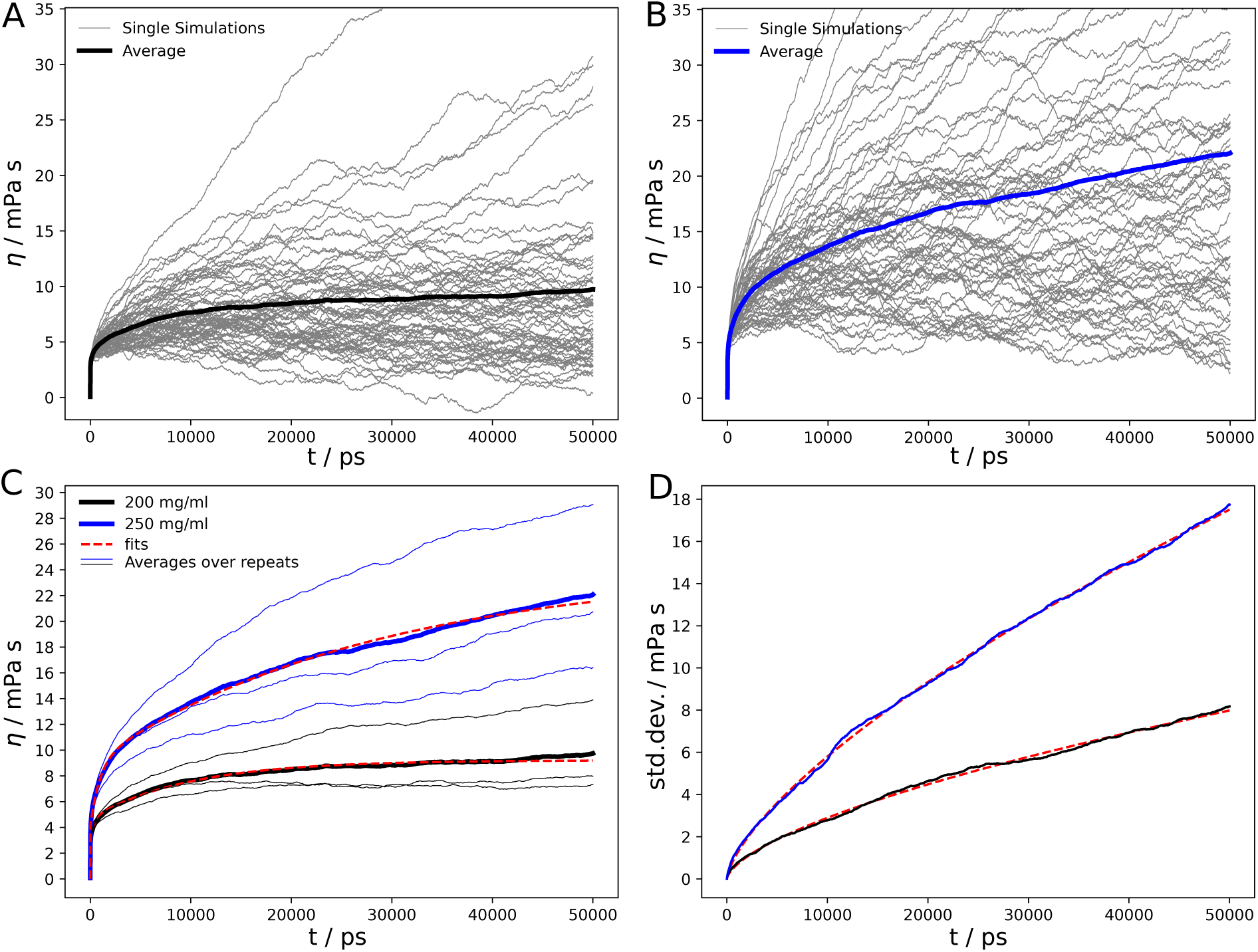
Green-Kubo integrals of the two mAb solutions studied. A) and B) Plotted are the running GK integrals of the *N* = 63 individual MD trajectories for the mAb concentrations of 200 mg/ml (A) and 250 mg/ml (B), together with the average (thick lines). C) The average GK integrals from A) and B) are plotted together with the triexponential fits (equation 4, dashed red lines). The averages over the three sets of repeats (each with *N* = 21 trajectories) are also plotted as thin lines. D) The standard deviations (equation 2) are plotted together with the fits (equation 3, dashed red lines).

**Table 1:** Shear viscosities (in mPa·s) of the mAb solutions determined with Eq. 4. The simulation sets 1, 2, and 3 refer to the sets of 21 MD trajectories initialized from each of the three 500-ns simulations. The statistical errors in the last column are the standard deviations of the three values from the three sets.

| mAb concentration | set 1 | set 2 | set 3 | combined |
| --- | --- | --- | --- | --- |
| 200 mg/ml | 7.5 | 14.4 | 7.2 | $9.2 \pm 5.8$ |
| 250 mg/ml | 31.2 | 20.3 | 24.4 | $24.1 \pm 7.8$ |

The slow rise in the running integrals (Figure 3C) show that for the mAb systems under study, there are long transients in the establishment of the asymptotic viscosities. These slow components are reflected in a large value of one of the three time constants in the fit (*τ_i_* in equation 4), which are 9.9 ns and 28.4 ns for *ρ* = 200 and 250 mg/ml, respectively; this slow component has a two times smaller amplitude at 200 mg/ml than at 250 mg/ml. Nevertheless, with the given sampling, the long-time viscosities can be determined with reasonable statistical precision, a notion that is also supported by the standard deviations (Figure 3D, see also bootstrap analysis below). We estimated the uncertainties in the final viscosities on the basis of the variation between the sub-averages over each of the three sets of simulations, each of which comprised of 21 MD trajectories of length 100 ns. Performing the triexponential fit to each of these three individual curves (which are also plotted in Figure 3C) yielded the values listed in Table 1.

The final viscosities obtained from the MD simulations are lower than the experimental viscosities of this mAb, which we determined with rotational rheometry (Figure 4 and Table 2). The computational viscosity of 24.8 ± 7.8 mPa·s at a mAb concentration of 250 mg/ml matches the experimental value of 23.2 mPa·s at 213 mg/ml, indicating different activity coefficients of the mAb solutions in the simulations and experiments (see the more detailed discussion of this aspect below).

**Figure 4:**
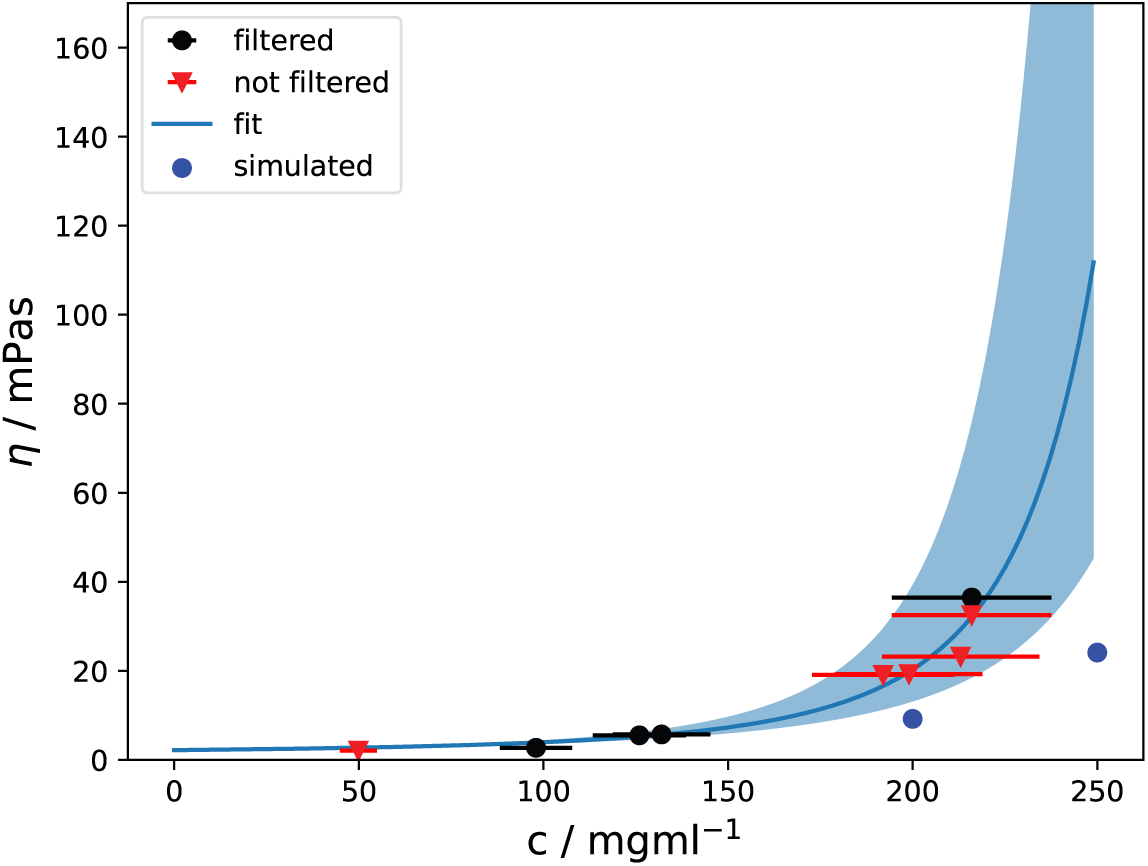
Experimentally determined dynamic viscosities at different mAb concentrations and Ross-Minton fit to the data. The black dots and orange triangles show the data from the filtered and unfiltered samples, respectively. The error bars reflect an uncertainty in the sample concentrations of ca. 10 %. The Ross-Minton fit to the combined data (filtered and unfiltered) is shown as a blue line. The error range, resulting from the min-max values of the concentration uncertainties, is depicted by the blue shaded area. The viscosities from the MD simulations are shown as blue dots.

**Table 2:** Dynamic viscosities of mAb solutions measured with rotational rheometry at 297 K.

| mAb concentration<br>(mg/ml) | viscosity<br>(mPa·s) |
| --- | --- |
| filtered |  |
| 98 | 2.7 |
| 126 | 5.5 |
| 132 | 5.7 |
| 216 | 36.5 |
| not filtered |  |
| 50 | 2.1 |
| 192 | 19.1 |
| 199 | 19.3 |
| 213 | 23.2 |
| 216 | 32.5 |

The data in Table 1 show a large difference between the viscosities obtained from the 3 independent sets of simulations at 200 mg/ml, with set 2 yielding a two times greater viscosity than sets 1 and 3 (a similarly large relative difference is not observed for the simulations at 250 mg/ml, with the viscosity from set 1 being only about 40% greater than the average from sets 2 and 3). A close inspection of the trajectories from the 200 mg/ml simulations revealed that in set 2 (but not in sets 1 and 3), more pronounced Fc-Fc domain contacts occur. The radial distribution functions (RDFs, Figure S3) show peaks at distances between the Fc domains of approximately 4 nm and 5 nm. Such peaks are not observed at 250 mg/ml (Figure S4). Figure S5 depicts one mAb dimer formed, with the Fc domains of the two mAbs in close proximity. These findings suggest that close Fc-Fc contacts could contribute to the high viscosity of mAb solutions. However, these apparent correlations should be interpreted cautiously, considering the statistical uncertainty. Nevertheless, previous observations have shown that the Fc regions contribute to solubility, and they have been utilized to improve solubility.^60–64^ Thus, mitigating Fc-Fc interactions, either via solvent design or via targeted mAb modifications through site-directed mutagenesis, might be a promising avenue toward less viscous and more stable mAb solutions.

To check the structural integrity of the mAbs during the course of the simulations, we analyzed the stability of the secondary structure elements over time and calculated the root-mean-square deviation from the starting structures of the simulations (Figure S6). No unfolding of structured parts of the Fab or Fc domains was observed in the simulations, and the overall structures of the individual mAb domains are stable during the simulations.

Concerning the nonuniform weighting of the data in the triexponential fits, the *b*-parameters were found to be 0.63 and 0.68 for the mAb concentrations of 200 and 250 mg/ml, respectively. Interestingly, these values are close to the *b* value of ca. 0.7 reported by Zhang et al. for a high-viscosity ionic liquid, for which their simulations yielded *η* = 19 mPa·s.^41^ In contrast, for the low-viscosity fluid ethanol, a *b*-value of 0.5 was found.^41^ This difference might be explained by the expected larger increase of *σ*(*t*) with time for more viscous fluids due to the sampling noise. Whether the match between the *b*-values found in our work for dense mAb solutions and the ones reported by Zhang et al. for ionic liquids, two systems that are on the one hand obviously very different but on the other both have high viscosity, are somewhat coincidental or might hint towards a common general underlying behavior remains an open question. In any case, simply assuming *b* = 0.5 instead of explicitly determining the exponent from the fit (equation 3) might not be a generally applicable approach.

To further quantify the statistical convergence of the computed viscosities, bootstrap analyses of subsets of trajectories were performed. In Figure 5 the viscosity estimates obtained from drawing random subsets comprised of a different number of MD trajectories are plotted, together with the standard deviations. The plots show that reasonable estimates can be obtained with ca. 40 trajectories.

**Figure 5:**
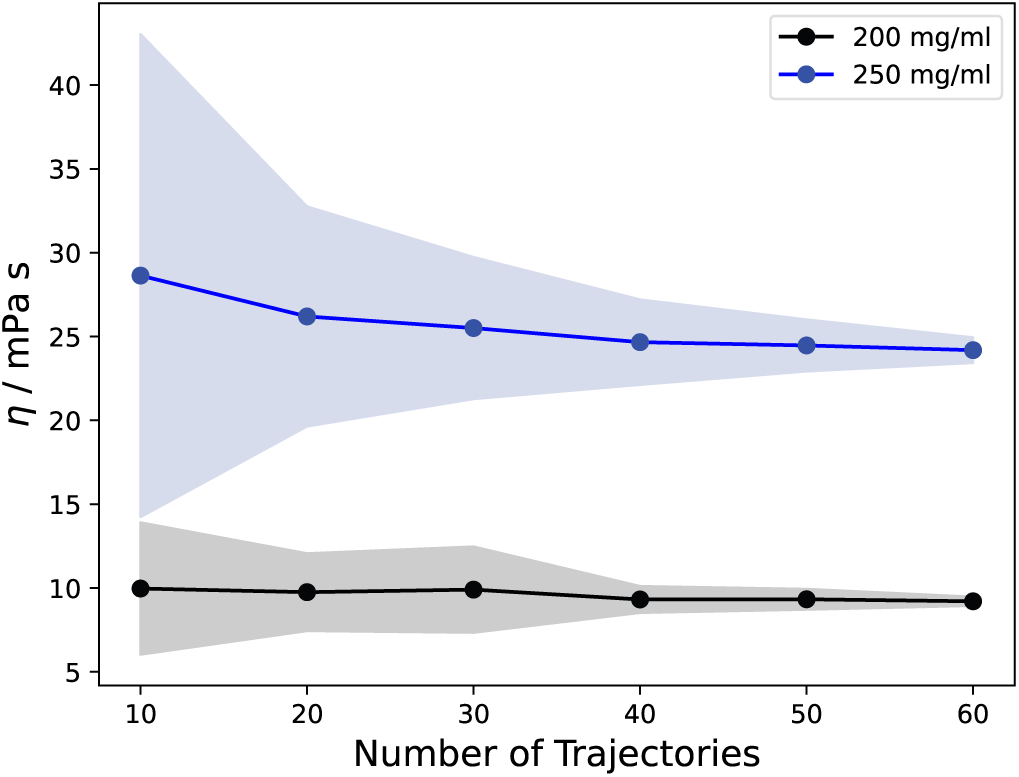
Bootstrap analysis of the statistical convergence of the viscosity estimate as a function of the number of MD trajectories. The black and blue lines represent the average viscosity obtained for 200 mg/ml and 250 mg/ml concentration by randomly drawing the indicated number of trajectories from the full set. Each draw was repeated 100 times; the shaded areas show the standard deviation.

Finally, we also used the GK method to determine the viscosity of bulk water as a reference, as described with the a99SB-disp water model used in our mAb simulations. The resulting water viscosity of 1.07 mPa·s (Figure S7) is slightly higher than the 1.02 mPa·s estimated by Qiu et al. from water diffusion constants.^65^ As a control, the bulk water viscosity simulations were repeated with 2 fs integration time steps (instead of 4 fs used in the other simulations, see Methods), yielding identical results within the statistical errors (Figure S7). The bulk water viscosity of the a99SB-disp model is about 20% higher than the experimental value, which is 0.896 mPa·s at 298 K.^66^

## Discussion

On the one hand, it is encouraging that the MD simulations predict relatively high viscosities for the two mAb solutions investigated, especially in light of the formidable challenges involved with computing collective properties of biomolecular solutions in the high-viscosity regime via all-atom MD. In particular, the simulations clearly distinguish the viscous behavior of the two different mAb concentrations studied, with the solution having a twofold greater viscosity at 250 mg/ml than at the lower concentration of 200 mg/ml. On the other hand, the simulations slightly underestimate the experimental viscosity, indicating that there is room for improvement. In the following, we shall thus discuss possible reasons for this discrepancy. We start with the computational perspective and then turn to experimental aspects.

There are two principal limitations of MD simulations that are ubiquitous and always need to be considered, the force field challenge and the sampling challenge. The force field is the potential energy function that describes the intra- and intermolecular interactions in the system, and on the basis of the assumption that the high viscosity observed for the mAb solution is governed by a transient and dynamic higher-order network of mAb molecules, an underestimation of the viscosity might hint towards too weak effective mAb-mAb interactions in this network. Explicitly addressing this question is not only computationally very demanding but also conceptually difficult, because in addition to the direct mAb-mAb interactions also water-mediated interactions contribute and thus the force field terms that describe the mAb-water interactions. In addition to the polypeptide chains, also the glycan parts of the mAbs can contribute to the interactions, either through direct glycan-glycan or glycan-mAb interactions or indirectly, for example by shielding hydrophobic surface patches.^60^ Concerning the combination of the AMBER a99SB-disp protein force field with the GLYCAM06j carbohydrate force field, more systematic studies would be desirable.

The result that the viscosity of the mAb solutions is underestimated in the simulations might be somewhat counter-intuitive on the basis of the slight overestimation of the viscosity of bulk water with the a99SB-disp water model used in this work (by about 20%, see above). However, the viscosities of the high-concentration mAb solutions are roughly one order of magnitude larger and are governed by the strength and dynamic response of the three-dimensional mAb network, not by the properties of the water itself.

The comparison of experimental and simulated viscosities is challenging because in the high-viscosity regime, the shear viscosity strongly increases in a nonlinear way with increasing mAb concentration. This means that even a small “shift” or “offset” in the concentration leads to a rather large viscosity change. In the context of the discussion of possible limitations of the MD force field, such a shift could result from a slight underestimation of the effective mAb-mAb interactions, resulting in a lower effective mAb concentration (or activity) in the simulations compared to the experiments. The computational viscosity at 250 mg/ml concentration matches the experimental one at a concentration of 213 mg/ml. Thus, if one would use viscosity as a gauge, these concentrations would correspond to a ratio of the computational and experimental mAb activity coefficients, 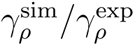, of ca. 0.8. A similar ratio is obtained by relating the MD-derived viscosity at 200 mg/ml to the mAb concentration obtained from the Ross-Minton fit for the corresponding viscosity value, which is ca. 165 mg/ml. (However, in general, 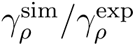 is expected to be concentration-dependent and approach 1 in the dilute limit.) The difference of ca. 20% found between the activity coefficients from the MD simulations and the experiments is relatively small and in our view reasonable given the many approximations made in the computational modeling. Due to the exponential increase of the viscosity with mAb concentration, the ratio of 0.8 translates into a large viscosity difference in the high-concentration regime.

The AMBER a99SB-disp force field^43^ used in the MD simulations was developed on the basis of the previous a99SB parameter set for providing a more balanced description of both disordered and folded globular proteins and, together with the associated water model that was derived from the original TIP4P-D water potential,^67^ to fix issues related to too attractive protein-protein interactions found in previous force fields.^68^ Of course, it could be possible that this reparametrization overshot the goal, in the sense that the effective mAb-mAb interactions are now too weak. However, there are several examples in the literature in support of the accuracy of this force field for protein-protein interactions, which for example govern the compactness of the structural ensembles of intrinsically disordered peptides or proteins. Notably, this was not only shown for relatively short disordered peptides^69^ but also for the more challenging case of large proteins with extended disordered regions.^70^ Taken together, we consider it to be unlikely that force field insufficiency alone is responsible for the lower viscosities in the simulations compared to the experiments. Other aspects have to be taken into consideration as well, in particular sampling limitations.

Viscosity is a measure for the dissipation of energy due to the deformation of a fluid. The underestimated viscosities in the simulations suggest that the energy dissipation is actually slower than predicted by the simulations, which presumably underestimate a slowly relaxing collective coupling mode between neighboring mAbs in the dynamic network. A basic question related to the MD simulations is whether the time scales sampled are long enough to capture the slow relaxation processes that are related to the reconfiguration of the dynamic network formed by the clusters of mAb molecules, in addition to finite size effects (see above). It is reasonable to assume that the time scales of these reconfiguration motions are governed by the diffusion dynamics of the mAbs. Riddiford and Jennings ^71^ obtained a rotational correlation time of 157 ns at 298 K for a bovine IgG from birefringence decay measurements, which is in line with the value of 168 ns determined by fluorescence anisotropy for a rabbit IgG.^72^ These values have been extrapolated to infinite dilution, and under high-concentration conditions a slowdown by a factor of 5 to 10 can be expected.^73^ The resulting time scales in the *µ*s range are in line with the broad signal in the 0.1 to 10 MHz frequency range (0.1 to 10 *µ*s) observed in dielectric relaxation spectroscopy of dense mAb solutions, which was assigned to originate from mAb reorientation motions.^57^ However, mAbs are very flexible molecules in which the individual Fab and Fc domains are connected via a flexible hinge and thus undergo substantial motions relative to each other. Hence, the rotational correlation time of the overall molecule might not be as relevant as the individual domain reorientations, which are faster. Indeed, Yguerabide et al. found that the Fab portions of the antibody are freely rotating over an angular range of ca. 30 degrees with a rotational correlation time of 33 ns.^72^ In any case, if such domain motions govern the dynamic viscosity they would need to be sampled in the MD simulations. These dynamics could be underlying the observed long transients in the GK integrals. The challenges of sampling such slow motions are notoriously hard, because in order to compute statistically reliable autocorrelation functions, the time scales of the simulations should exceed the desired relevant correlation times by at least a factor of 50, preferably even more. In the simulation protocol applied in this work, this is implemented via averaging over the *N* = 63 MD trajectories generated for each system. The time scale sampled in each individual trajectory is 100 ns, and hence the onset of the domain reorientation motions can be sampled. From the rise of the GK integrals and the corresponding slow components in the triexponential fits, we conclude that parts of these slow dynamics are sampled in the MD simulations (see Figure S8 for an analysis of the orientational diffusion of the individual Fab and Fc domains). This conclusion is further supported by an additional check, in which only the first 20 ns of the running integrals were used for the triexponential fits (and hence only dynamics up to that time scale contribute). These fits yielded long-time viscosities of 8.5 and 18.3 mPa·s for 200 and 250 mg/ml, respectively, which are reasonably similar (within the statistical errors) to the respective values from the full GK integrals (Table 1).

After having discussed potential limitations on the computational side, also experimental issues linked to viscosity measurements of dense mAb solutions will be considered. First, the dynamic viscosity should be a constant (independent of stress), that is, the solution should behave as a Newtonian fluid. In practice, however, the measured viscosities might depend on the shear rate, which led to the convention to report rotational rheometry data at a shear rate of 1000 s*^−^*^1^. When comparing experimental viscosities with computational values, one should keep in mind that the GK approach yields zero-shear viscosities. But even if the mAb solution behaves reasonably Newtonian, additional challenges remain. For example, as shown and discussed by Pathak et al., adsorption of mAbs at the air/water interface can affect conformational stability and foster the formation of irreversible particles, which can lead to a substantially increased apparent viscosity.^74^ This overestimation was shown to be less severe for double gap rheometers compared to cone plate ones, such as the rheometer used in the present work.^74^ Removing the largest particles via filtration of the solution before the viscosity measurement diminishes the effects, but does not entirely eliminate them. As described above, in the present study rotational rheometry measurements were done with both unfiltered and filtered samples and no systematic differences were recorded.

## Conclusions

The accurate computational prediction of collective properties of dense biomolecular solutions, such as the viscosity, is challenging but highly desirable, not only for studies of the crowded interior of biological cells or the formation of biomolecular condensates but also for the development of high-concentration biopharmaceutical formulations. Experimental characterization of high-concentration biopharmaceutical formulations is time- and material-consuming and thus difficult, especially in the early development phases where material is limited.

The present work is a proof of concept study that demonstrates the use of all-atom MD simulations for the computational prediction of the dynamic viscosity of high-concentration monoclonal antibody solutions from the pressure fluctuations using the Green-Kubo ap-proach. The Padlan IgG1 mAb was used as a model system. Large-scale MD simulations of systems representing mAb concentrations of 200 and 250 mg/ml yielded viscosities of 9.2 and 24.1 mPa·s, respectively, demonstrating that assessing the challenging high-viscosity regime is indeed feasible using MD simulations. However, the experimental viscosities of this mAb are 19 and 23 mPa·s at 199 and 213 mg/ml, respectively, showing that there is still room for improvements.

The underestimation of the viscosity in the MD simulations might hint at slightly too weak effective interactions between the investigated IgG1 mAbs in the AMBER a99SB-disp force field that could result in a too loose dynamic mAb network, which is the microscopic origin of the high viscosity. However, this interpretation should not be overgeneralized, because a larger and more diverse set of dense protein solutions would need to be investigated. Such a systematic study would be computationally very demanding, and the present study highlights the immense sampling challenges involved with computing the viscosity of dense biomolecular solutions. However, at the same time, it demonstrates the principal feasibility of such an endeavor. Thereby, it also opens the way towards using all-atom MD simulations to investigate in microscopic detail the modulation of the viscosity of mAb formulations, for example by site-directed mutagenesis^75–80^ or by solvent design, for example via the addition of excipients.^81–85^ In that context, it appears promising to synergetically combine rigorous but computationally expensive physics-based methods, such as all-atom MD simulations, with efficient heuristic approaches.^86,87^ From a more general perspective, the finding that MD simulations can describe the high-viscosity regime of dense protein solutions also has implications for atomistic simulations of other crowded environments, such as biomolecular condensates and the cytoplasm.

## Data and Software Availability

All relevant data are included in the Tables and Figures provided in the manuscript and Supporting Information. MD simulations were performed with GROMACS version 2020.5 (https://manual.gromacs.org/documentation/2020.5/download.html). The code for the viscosity calculations is available on github (https://github.com/TobiasMPrass/gmx_gk_autocorr). The MD parameter file, starting coordinates, and force field topology files are enclosed as Supporting Information.

## Supporting Information Available

Comparison of double- and triple-exponential fits of the GK integrals. Comparison of the normalized pressure ACF of bulk AMBER a99SB-disp water to those of the concentrated mAb solutions. Description of the viscosity calculations of bulk water. RDF plots from the simulations at 200 and 250 mg/ml. Electrostatic potential of the Padlan mAb mapped onto the van-der-Waals surface and snapshot of mAb dimer from 200 mg/ml simulation set 2 showing close Fc-Fc contacts. Plots of C*α*-RMSD distributions and time evolution of secondary structure count during simulations. Plots of the bulk water GK integrals. Time-correlation functions showing reorientation motions (orientational diffusion) of the mAb domains.

## Acknowledgement

We thank Sören von Bülow for useful discussions, Moritz Schmidt, Sebastian Andris, Tobias Hepp, and Andrea Eiperle for technical support, and Raphael Drerup for project administration support. This work was supported by Boehringer Ingelheim Pharma GmbH & Co. KG and by Deutsche Forschungsgemeinschaft (DFG) under Germany’s Excellence Strategy EXC 2033 - 390677874 - RESOLV. The authors gratefully acknowledge the funding of this project by computing time provided by the Paderborn Center for Parallel Computing (PC2).

## Supporting Information

### Double-versus tripe-exponential fits

**Figure S1:**
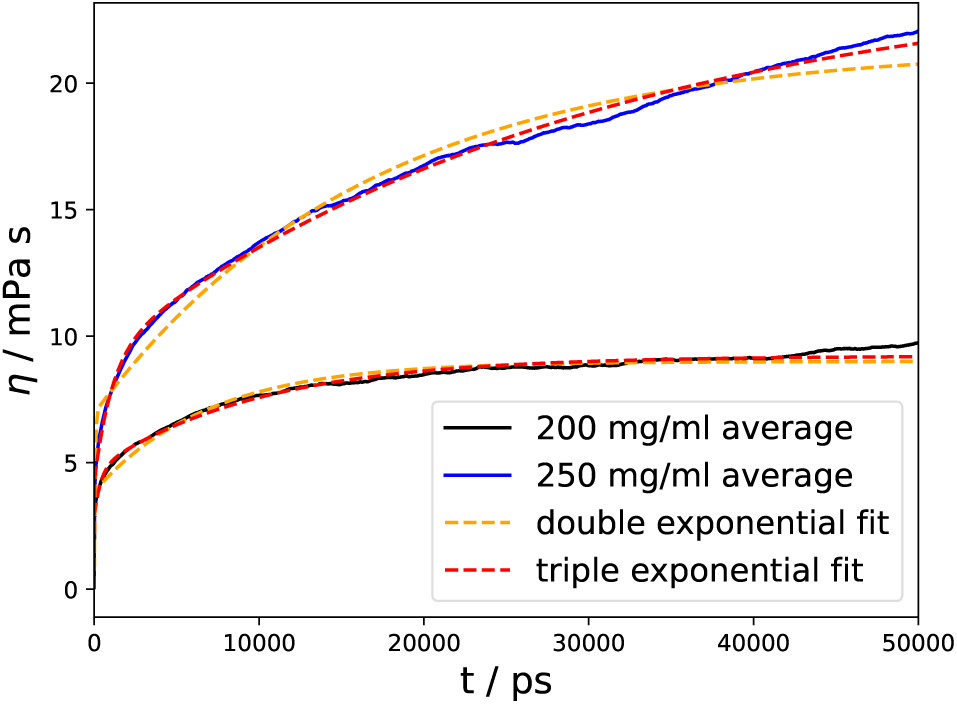
Comparison of a double- and triple-exponential fits to the viscosity curves of the concentrated mAb solutions. The triple-exponential fits, depicted with red dashed lines, capture the actual data at both 200 mg/ml (black curve) and 250 mg/ml (blue curve) more closely than the double-exponential fits (orange dashed lines; the long-time limiting values of the viscosities from the double-exponential fits are *η* = 9.0 and *η* = 21.5 mPa·s for 200 and 250 mg/ml, respectively).

### Viscosity of bulk AMBER a99SB-disp water model

**Figure S2:**
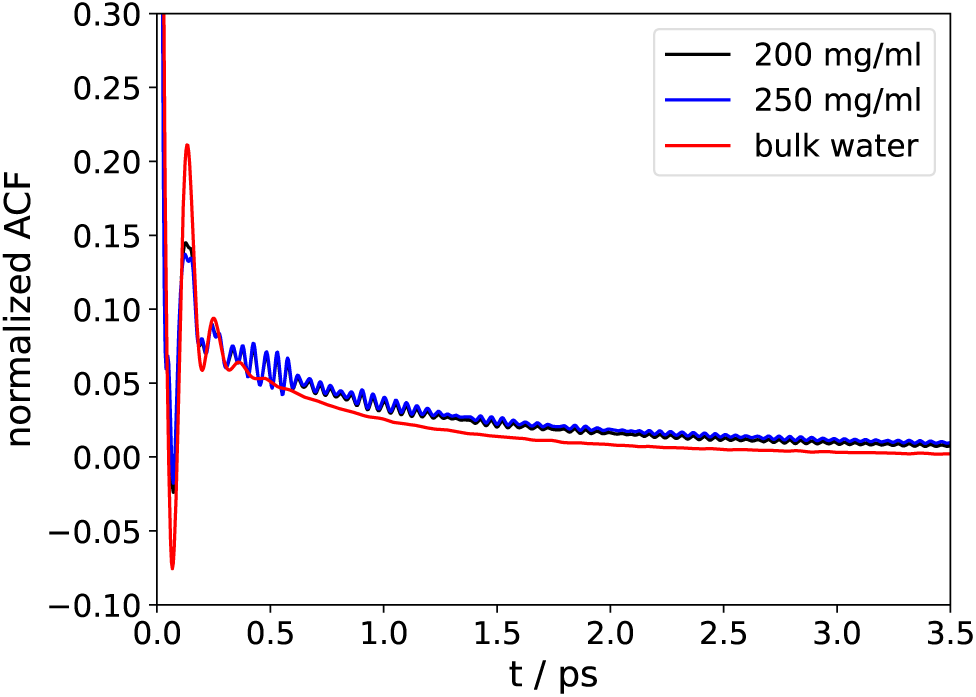
Comparison of the normalized pressure autocorrelation functions (ACFs) between the antibody systems and bulk water (a99SB-disp water model^S1^).

To determine the shear viscosity of bulk a99SB-disp water, a cubic box of 2000 water molecules was simulated in the NpT ensemble at 298 K for 500 ns. Similar to the mAb simulations, 21 configurations taken from the last 400 ns were used to initiate 5 ns NVT simulations during which the pressure was saved to disk every 4 fs. To compute the shear viscosity from the GK integrals, a similar procedure as used for the mAb solutions was followed, with the exception that a double-exponential fit (equation 1)

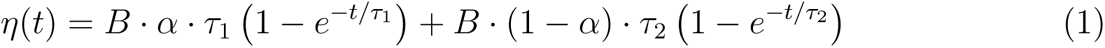

with fitting parameters *B >* 0, *α, β <* 1, and *τ*_1,2_ *>* 0, was used to extrapolate the GK integrals to the long time limit.

The 21 viscosity graphs and average are shown in Figure S7. The viscosity of bulk a99SB-disp water obtained from the fit to the average of the 21 GK runnning integrals is 1.07 mPa·s, which is slightly higher than the value of 1.02 mPa· reported by Qiu et al. (2021),^S2^ who estimated the viscosity from the self-diffusion coefficient. The a99SB-*disp* water model thus overestimates the experimental value of 0.896 mPa·s at 25 *^◦^*C^S3^ by ca. 20%.

**Figure S3:**
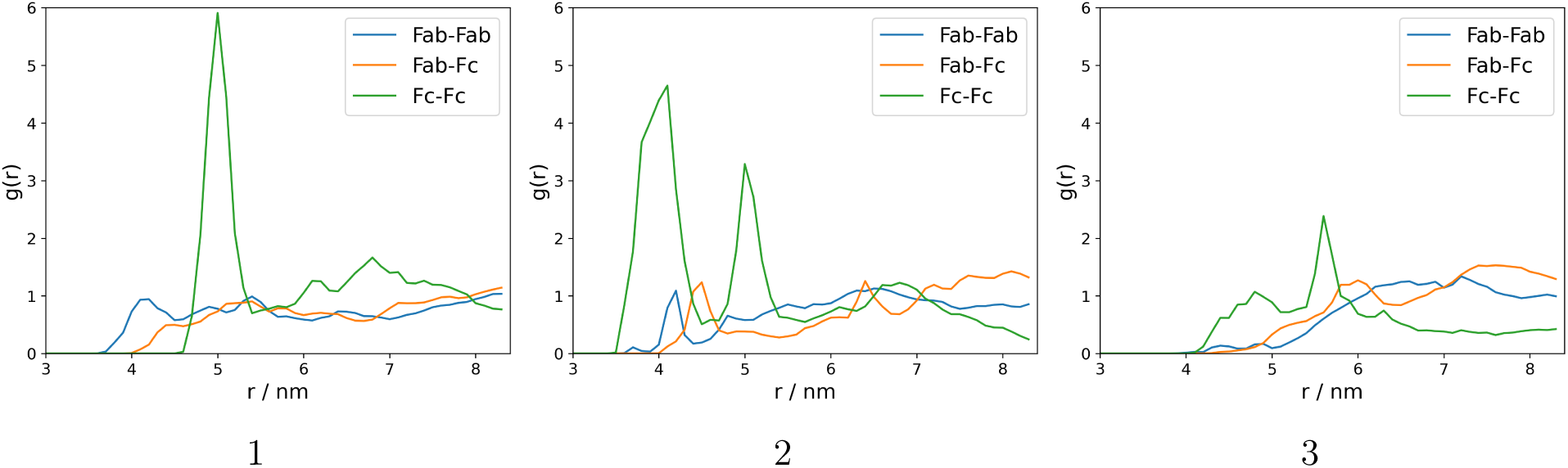
Radial distribution functions of the antibody domains with respect to the domain centers-of-mass in the 200 mg/ml simulation sets 1, 2 and 3.

**Figure S4:**
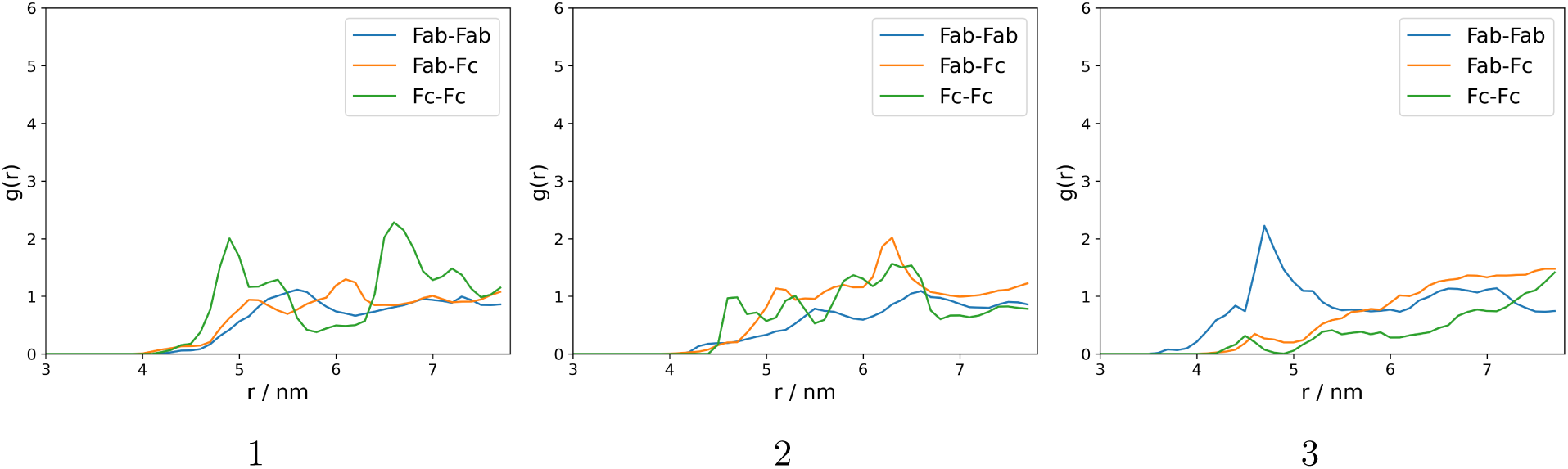
Radial distribution functions of the antibody domains with respect to the domain centers-of-mass in the 250 mg/ml simulation sets 1, 2 and 3.

**Figure S5:**
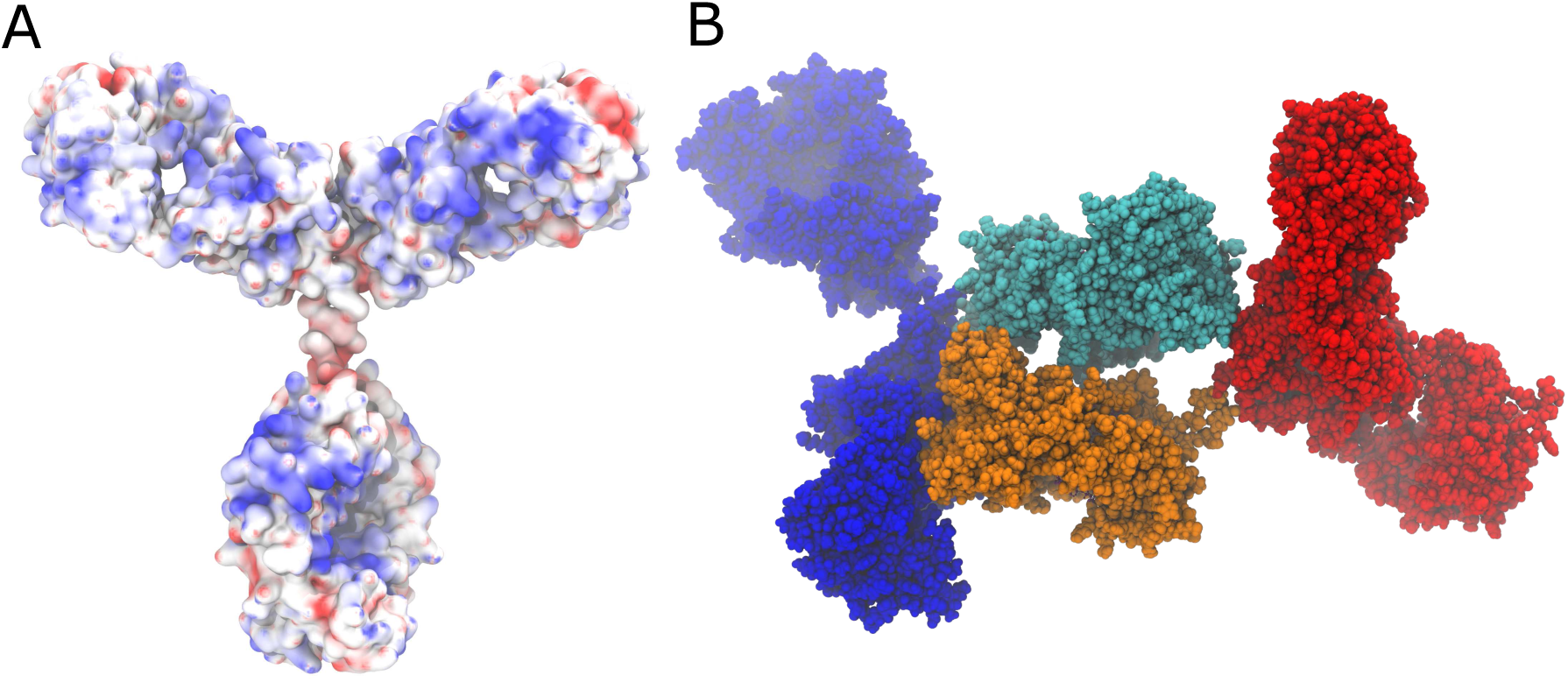
A) Visualization of the electrostatic potential mapped onto the mAb surface. B) Snapshot taken from “set 2” of the 200 mg/ml simulations, showing 2 of the mAbs forming a dimer via the Fc domains (colored in cyan and orange).

**Figure S6:**
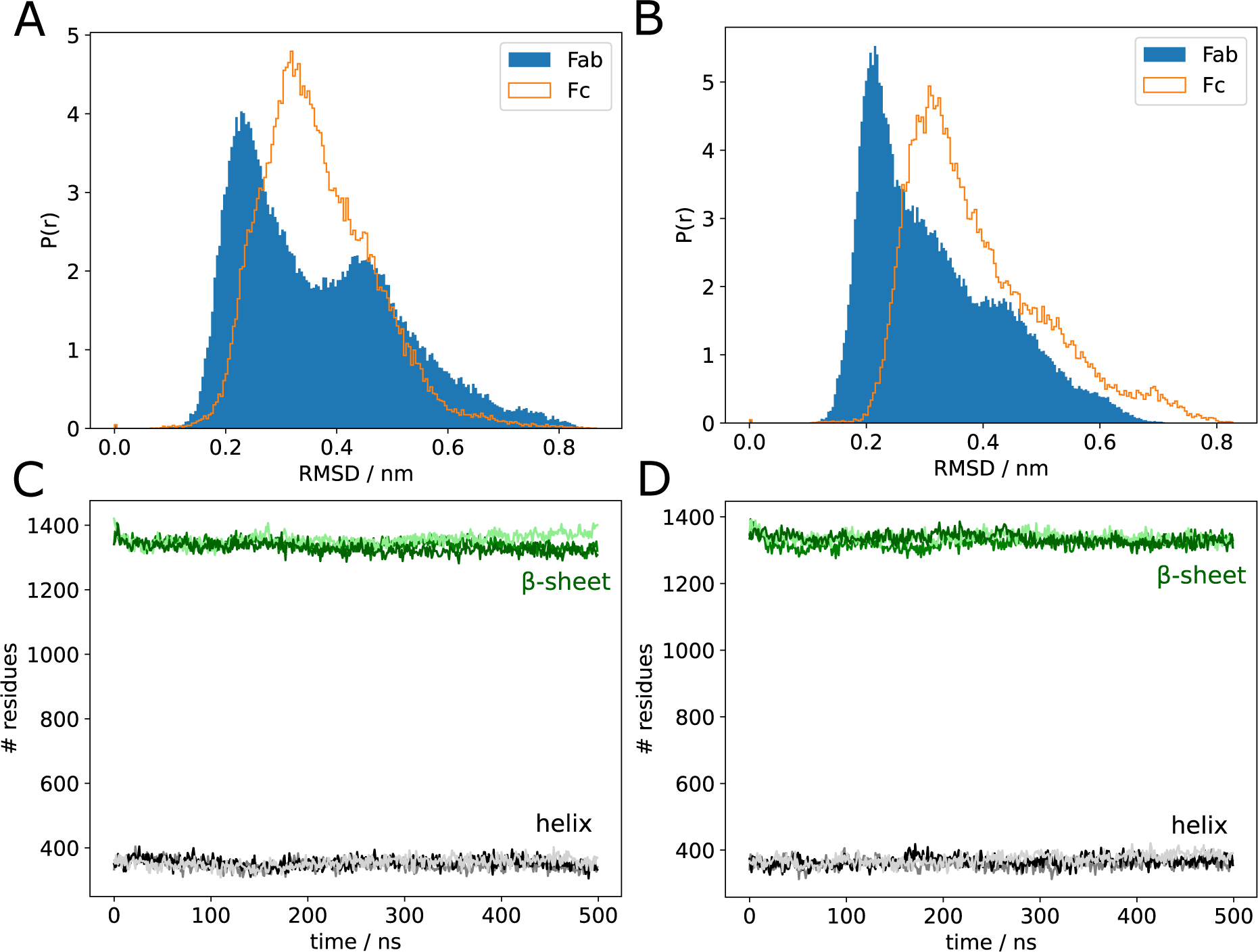
Analysis of the structural integrity of the mAbs in the simulations. A) and B) show the C*α*-RMSD distributions in the 200 mg/ml and 250 mg/ml mAb solutions, respectively. C) and D) show the time evolution of the number of residues in *β*-sheets and helices in the 200 mg/ml and 250 mg/ml systems, respectively. This residue count is over all 4 mAbs in the simulation systems.

**Figure S7:**
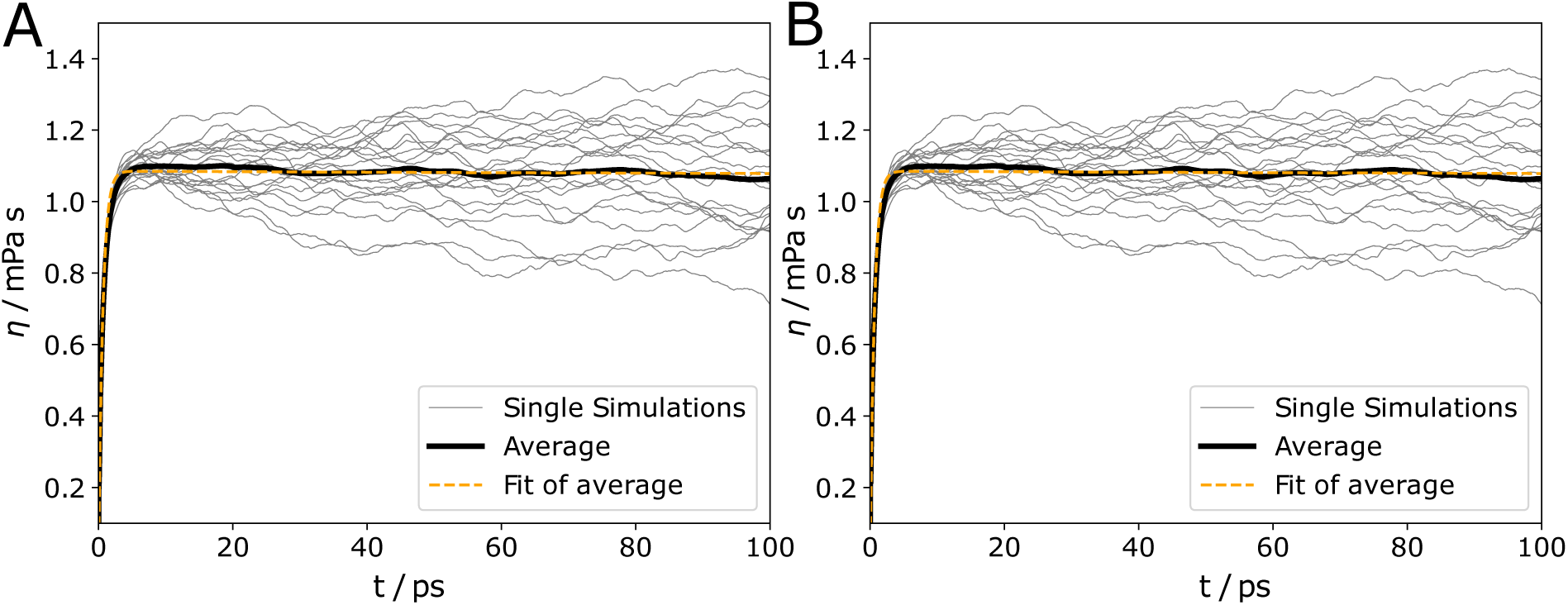
Running GK integrals from 21 individual MD trajectories of bulk water (a99SB-disp water model), computed in with integration time steps of A) 2 fs and B) 4 fs. The individual integrals are plotted as thin grey lines, the average is plotted as a thick black line. The dashed lines show the double-exponential fits, 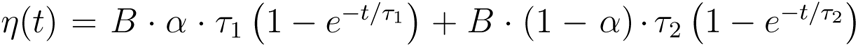, which yield long-time limit viscosities of *η* = 1.0757 and 1.0745 mPa·s for the simulations with the 2 fs and 4 fs integration time steps, respectively.

**Figure S8:**
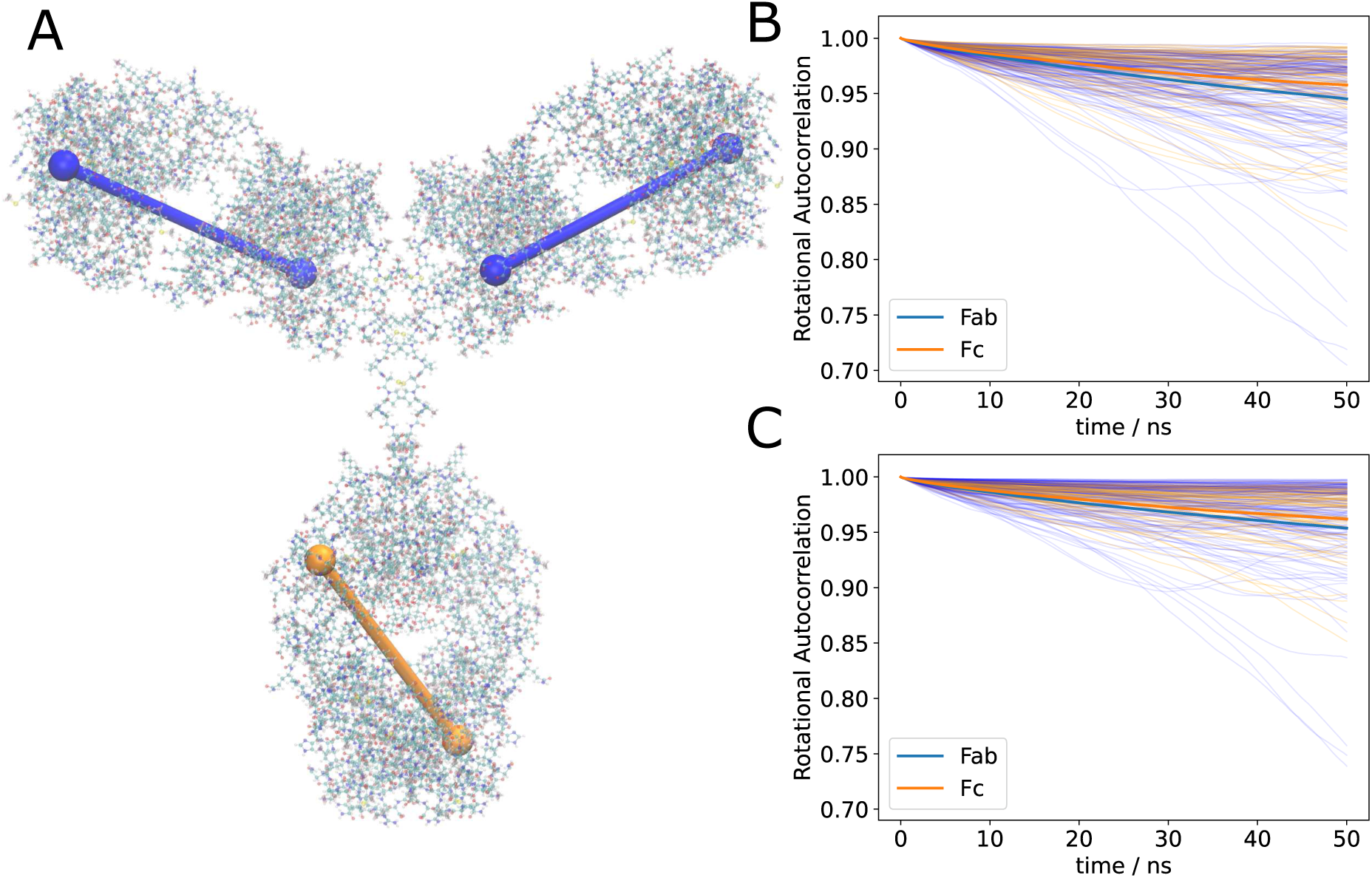
Time correlation functions (TCFs) describing the reorientation motions (orientational diffusion) of the mAb domains. Here, the TCF of the cosine of the angle *α* is plotted, where *α* is the angle between a vector at time *t*_0_ and at time *t*_0_ + *t*. A) shows the analyzed vectors that span across the individual mAb domains (Fab: LC Ala72 to HC Cys209 (blue); Fc: HC1 Trp286 to HC2 Val357 (orange)). B) and C) show the TCFs of the Fab (blue) and Fc (orange) domains in the 200 mg/ml and 250 mg/ml systems, respectively.

**Figure S9:**
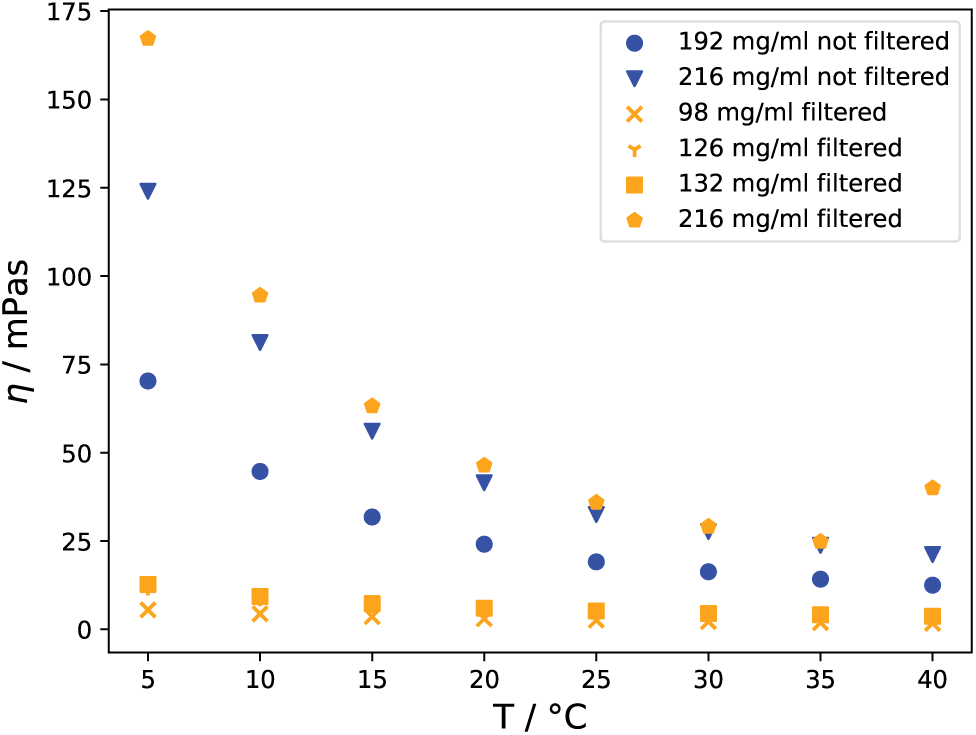
Rotational rheometry viscosities from the temperature ramps. The data from the filtered and unfiltered samples are depicted in orange and blue, respectively.

## Notes

https://github.com/TobiasMPrass/gmx_gk_autocorr

### Competing Interest Statement

The authors have declared no competing interest.

